# Selective attention and decision-making have separable neural bases in space and time

**DOI:** 10.1101/2021.02.28.433294

**Authors:** Denise Moerel, Anina N. Rich, Alexandra Woolgar

**Affiliations:** School of Psychological Sciences, Macquarie University, Sydney, Australia; Perception in Action Research Centre, Macquarie University, Sydney, Australia; Macquarie University Performance and Expertise Research Centre, Sydney, Australia; MRC Cognition and Brain Sciences Unit, University of Cambridge, Cambridge, United Kingdom

## Abstract

Attention and decision-making processes are fundamental to cognition. However, they are usually experimentally confounded, making it difficult to link neural observations to specific processes. Here we separated the effects of selective attention from the effects of decision-making on human brain activity, using a two-stage task where the attended stimulus and decision were orthogonal and separated in time. Multivariate pattern analyses of multimodal neuroimaging data revealed the dynamics of perceptual and decision-related information coding through time (magnetoencephalography (MEG)), space (functional Magnetic Resonance Imaging (fMRI)), and their combination (MEG-fMRI fusion). Our MEG results showed an effect of attention before decision-making could begin, and fMRI results showed an attention effect in early visual and frontoparietal regions. Model-based MEG-fMRI fusion suggested that attention boosted stimulus information in frontoparietal and early visual regions before decision-making was possible. Together, our results suggest that attention affects neural stimulus representations in frontoparietal regions independent of decision-making.

## 1. INTRODUCTION

Selective attention is a mechanism that prioritises relevant information from amongst competing sensory input. With clear behavioural benefits to attending to particular locations in space (Pestilli & Carrasco, 2005; Posner, 1980) or visual features (Sàenz et al., 2003; White & Carrasco, 2011), there is strong evidence that attention affects perception (Carrasco, 2011). However, in cognitive neuroscience experiments, attended information is also commonly the information that the participant makes a decision about, confounding the processes of selecting and maintaining relevant stimulus information with decision-making processes. In this study, we separate the effects of attention on the neural processing of information from the effects of decision-making and investigate the spatiotemporal correlates of each process separately.

According to the adaptive coding hypothesis (Duncan, 2001), single cells in frontoparietal cortex flexibly adapt their response to code different information as needed for the task, providing a potential source of bias for more selective brain regions (Desimone & Duncan, 1995). In humans, a network of frontoparietal regions, which we will refer to as the ‘Multiple Demand’ (MD) regions (Duncan, 2010; Duncan et al., 2020; Duncan & Owen, 2000), responds with a profile consistent with adaptive coding. Functional Magnetic Resonance Imaging (fMRI) studies in human participants show that the MD regions respond to a wide range of cognitively demanding tasks (Assem et al., 2020; Dosenbach et al., 2006; Duncan & Owen, 2000; Fedorenko et al., 2013).

More specific evidence for an adaptive response in MD regions comes from fMRI studies using multivariate pattern analyses (MVPA). These studies show that the MD regions code for a range of task-relevant information in different contexts (Woolgar et al., 2016). In addition, several studies have found stronger coding in the MD cortex for cued (i.e., attended, task-relevant) objects (Woolgar, Williams, et al., 2015) or features (Jackson et al., 2016; Jackson & Woolgar, 2018) relative to equivalent distractors. In parallel, recent magnetoencephalography (MEG) and electroencephalography (EEG) studies have shown the coding of a cued feature at a cued location is sustained over time, while information coding for distractor information is not sustained (Barnes et al., 2022; Battistoni et al., 2020; Goddard et al., 2022; Grootswagers et al., 2021; Kaiser et al., 2016; Moerel et al., 2022; Noah et al., 2023). What we do not know, however, is to what extent this difference in information coding between cued and distractor stimuli is driven by decision-making processes pertaining to the cued information.

Underscoring the importance of this confound, there is clear evidence that decisions can drive decoding in early visual areas (Löffler et al., 2019; Rens et al., 2017), and frontoparietal cortex (Löffler et al., 2019), although these studies do not explicitly investigate the role of attention, and the decision was not driven by visual information. Bode and colleagues (2012) found perceptual decisions could be decoded from 140-180ms post-stimulus onwards from pure noise images, slightly earlier than feature-based attention effects (Goddard et al., 2022). Thus, decisions can be decoded in similar brain areas, with a time-course that could interact with that of attention effects, further complicating the interpretation of the extant literature. There is some evidence suggesting that attention effects can occur in the absence of decision-making (Hon et al., 2006). However, this study did not investigate what information was coded about the attended and unattended stimuli.

Here, we used MVPA of MEG and fMRI data to address 3 research questions. First, we used MEG to ask whether attention affects the coding of stimulus information when it is separated from decision-making processes in time, using a 2-stage task to dissociate the coding of attended and unattended visual information from decision-related information. Second, we used fMRI to ask what type of information the MD regions hold. Finally, we used model-based MEG-fMRI to formally combine the data from the two imaging modalities and examine the time-course with which the MD regions preferentially coded for cued compared to distractor information during the first phase of the trial.

## 2. MATERIALS AND METHODS

### 2.1. MEG acquisition and analysis

#### 2.1.1. Participants

The first part of this study consisted of a behavioural training session and a MEG session carried out at Macquarie University (Sydney, Australia). We tested 31 healthy volunteers in the behavioural training session. Of these, 21 performed well enough to participate in the MEG session (at least 90% accuracy in the final run of the training session). The MEG data from 1 participant were excluded due to a failure to complete the experiment, resulting in a final MEG sample of 20 participants (14 female/6 male, 18 righthanded/2 lefthanded, mean age = 25.8 years, SD = 5.1). Participants received $15 AUD for participation in the training session and $30 AUD for participation in the MEG session. The study was approved by the Macquarie University Human Research Ethics Committee, and all participants provided informed consent.

#### 2.1.2. Stimuli and experimental procedure

##### MEG session

Participants were instructed keep fixation at a central bullseye throughout the experiment and were cued to attend to either blue or orange at the start of each block (32 trials). Figure 1 shows the stimuli and trial procedure. We used a 2-stage task, to be able to separate the effect of attention from the decision-making process in time. In stage 1, approximately equiluminant blue and orange oriented lines within a circular window were overlaid at fixation for 150ms on a mid-grey background (Figure 1A). Participants attended to the lines of the cued colour, while ignoring the lines of the other colour. After a 500ms blank screen, a black comparison line was presented for 200ms (stage 2). The task was to determine which way the cued lines had to be rotated to match the orientation of the comparison line. After the comparison line was presented, there was a 500ms blank screen, and then the response screen, consisting of arrows showing either a clockwise or an anti-clockwise rotation, was shown until a response was given. If participants did not respond within 3 seconds, the next trial started. Participants pressed one of two buttons to indicate whether the rotation shown on the response screen was correct or incorrect. We used this response screen to separate the correct response button from the rotation direction decision, ensuring that rotation direction decoding could not be driven by motor preparation. The mapping of the buttons to indicate ‘correct’ or ‘incorrect’ was counterbalanced across participants. Participants received feedback on every trial; at the end of each trial, the white part of the fixation bullseye turned green (when correct) or red (when incorrect) for 250ms to give feedback on accuracy.

**Figure 1.**
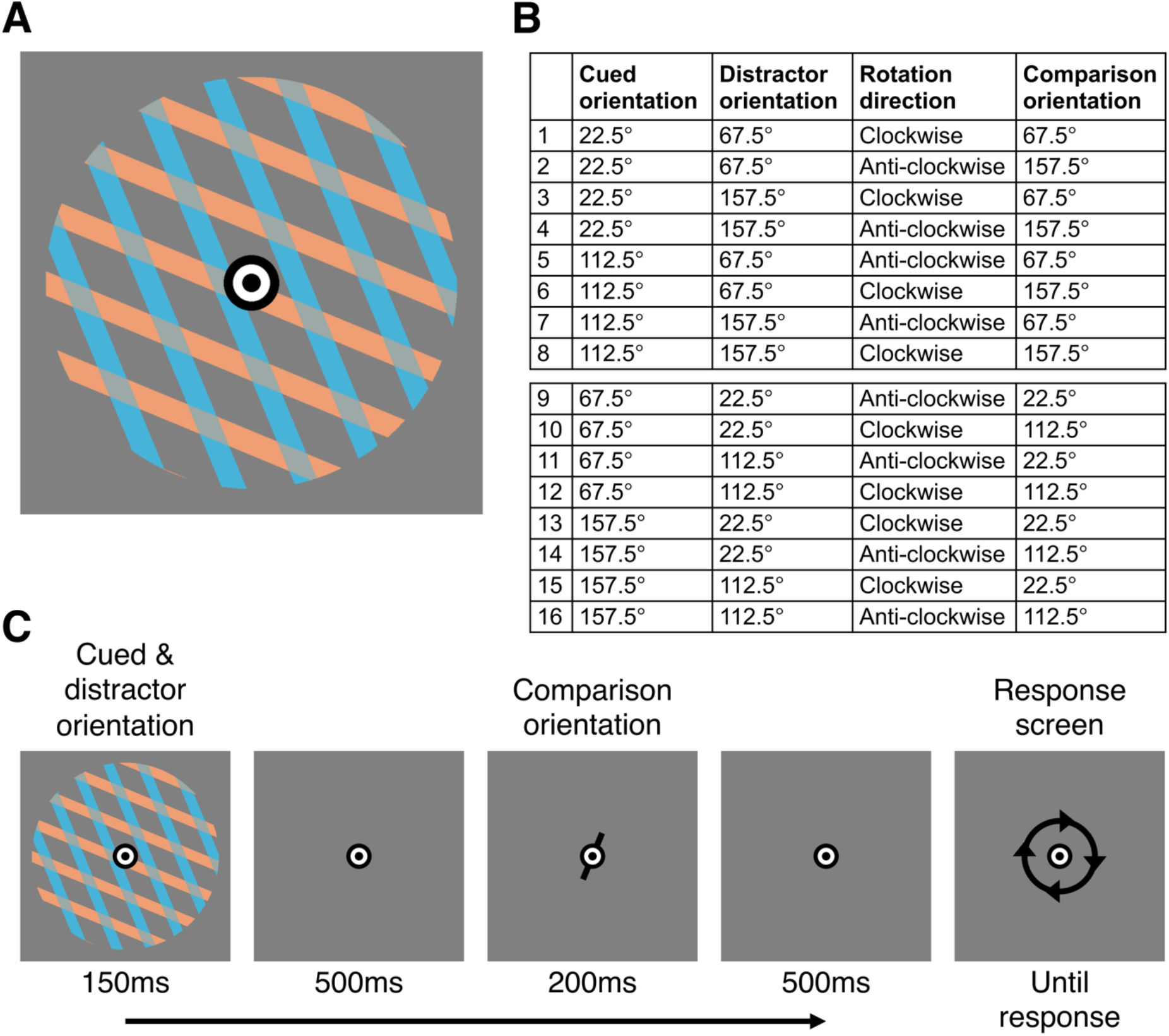
Stimuli and design. The stimulus, shown in **A**, consisted of oriented blue and orange lines within a circular window. Participants attended to the lines of one colour, while ignoring the lines of the other colour. **B** shows the 16 possible trial types, determined by a combination of cued orientation, distractor orientation, and rotation direction. The comparison orientation was not used in the analysis but is shown for completeness. The cued orientation, distractor orientation, and rotation direction are orthogonal dimensions in this design, only within the 2 pairs separately (top half and bottom half of the table). Decoding was done within each pair to ensure all of these factors are balanced. **C** shows an example of a trial sequence. Participants were cued to a particular colour for a block of 32 trials. On each trial, the cued and distractor oriented lines were shown for 150ms followed by 500ms delay, and then the comparison orientation was shown for 200ms. After another 500ms delay, the response screen appeared until a response (‘correct’ vs ‘incorrect’) was given, or until the 3s timeout. In this trial, for example, if the cued colour was blue, the participant would have to decide whether the blue lines need to rotate clockwise or anti-clockwise to match the comparison orientation. At the response screen, which in this trial shows clockwise, the correct response would be the button indicating ‘correct’.

The coloured oriented lines had a circular window of 3 degrees of visual angle (DVA), a spatial frequency of 2 cycles/degree, were 2 possible colours (blue; RGB = 239, 159, 115 and orange; RGB = 72, 179, 217), and were phase randomised (Figure 1A). The comparison line had a length of 0.8 DVA and a width of 0.1 DVA. The response mapping arrows were 1.5 DVA, and the central fixation bullseye had a diameter of 0.4 DVA. There were 4 possible orientations for the cued, distractor, and comparison orientations, divided into 2 pairs: 22.5° was paired with 112.5°, and 67.5° was paired with 157.5° (Figure 1B). When the cued orientation was from one pair, the distractor and comparison orientation were always from the other pair. This ensured that the two overlaid orientations were never the same. The comparison orientation was always rotated 45° from the cued orientation, either in a clockwise or an anti-clockwise direction.

Cued orientation (4) x distractor orientation (2) x rotation direction (2) x response screen (2) were counterbalanced within each block, and the cue colour was counterbalanced over blocks. Participants were instructed which colour to attend at the start of each block of 32 trials, and the cue colour switched between blocks. Participants completed 16 experimental runs, with 2 blocks per run (one block of 32 trials per cue colour). One participant completed 14 instead of 16 experimental runs. In addition to the feedback on every trial, participants received feedback on their accuracy at the end of each block. The order of the trial types was randomised within each block. To increase the number of correct trials, any trial on which the participant made an error was presented again later in the block. This was done a maximum of two times per block, and only if it was not the last trial in the block. We replaced error trials with successful retakes in the analysis, thus increasing the number of correct trials available for the decoding analysis. We did not record eye movements, but to influence decoding accuracy, eye-movements during the trials would have to systematically vary between the different conditions within each pair (see Figure 1B). This seems unlikely as all cued and distractor orientations were presented at fixation with no variance between conditions.

##### Training session

Due to the challenging task, participants completed a separate behavioural training session before doing the task in the MEG. The training session consisted of 3 parts. First, participants practiced the rotation task without selection. Only lines of the cued colour were shown, and the task was slower: the cued stimulus and comparison line were both presented for 500ms instead of 150ms and 200ms respectively. Next, the selection element was introduced. Both the cued and the distractor colour lines were shown, with the timing of the task again slowed as in part 1 of the training. Finally, the task was presented at the same speed as used in the MEG session: the cued stimulus was shown for 150ms and the comparison line was presented for 200ms. Feedback was given at the end of each trial by the white part of the fixation bullseye turning green (correct) or red (incorrect) for 250ms. At the end of each block, participants also received feedback about their overall accuracy for that block.

Each run within the training session consisted of 2 blocks of 32 trials, one block per cued colour. The number of runs per training part was adjusted flexibly depending on task performance. When participants had finished at least 2 runs and achieved an average accuracy of at least 90% for the previous run, they moved onto the next part of training. Participants who were able to complete all 3 training steps within an hour-long session were invited to participate in the MEG session.

#### 2.1.3. MEG acquisition

Participants performed the task in a supine position. The MEG signal was continuously sampled at 1000Hz on a whole-head MEG system with 160 axial gradiometers inside a magnetically shielded room (Model PQ1160R-N2, KIT, Kanazawa, Japan). All MEG recordings were obtained at the KIT-Macquarie Brain Research Laboratory, National Imaging Facility, Sydney, Australia. Recordings were filtered online between 0.03Hz and 200Hz. To track head movements during the experiment, participants were fitted with a cap with 5 marker coils. We recorded the head shape of participants using a Polhemus Fastrak digitiser pen (Colchester, USA). The stimuli were projected onto the ceiling of the magnetically shielded room. To correct for delays in stimulus onsets of the projector compared to the triggers, photodiode responses were used to determine the onset of the trial. We adjusted the stimuli to correct for distortions introduced by the projector. The stimuli were presented using the Psychtoolbox extension in MATLAB (Brainard, 1997; Kleiner et al., 2007; Pelli, 1997). Participants used a 2-button response pad to respond on each trial.

#### 2.1.4. Pre-processing

The data were pre-processed using the FieldTrip extension in MATLAB (Oostenveld et al., 2011). The recordings were sliced into 3100ms epochs; from 100ms before the onset of the first stimulus until 3000ms after stimulus onset. The signal was then down sampled to 200Hz. Error trials that had a successful retake were exchanged for the correct retake. To keep the counterbalancing of all conditions intact, we included the few error trials that did not have a successful retake in the analysis (0.33% of trials on average).

#### 2.1.5. Decoding analysis

We used MVPA to determine whether there was information present in the pattern of activation for each timepoint about: 1) the cued orientation; 2) the distractor orientation; and 3) the rotation direction. Our 2-stage task design allowed the effect of attention to be assessed from the first stage onwards, whereas the decision can only be made from the second stage onwards. This allows us to separate the effect of attention and decision-making in time. In stage 1 of the task, the cued and distractor orientation were shown. Comparing the coding of the cued and distractor orientations allowed us to determine whether attended visual information was preferentially coded. In stage 2, the comparison stimulus was shown. The coding of the rotation direction reflected the decision participants had to make about the stimulus. Critically, the cued orientation, distractor orientation, and rotation direction were all orthogonal dimensions, allowing us to investigate the coding of each of these separately. For each timepoint, we trained a Support Vector Machine (SVM) classifier on the activation across all 160 sensor channels to distinguish the conditions of interest. Classifier performance was determined using leave-one-run-out cross-validation. Data were split into a training dataset, containing the data from 15 runs, and a testing dataset, containing the data from the left-out run. This was done 16 times, using a different testing run each time. The analysis was repeated for each point in time, resulting in a decoding accuracy over time. We used the CoSMoMVPA toolbox for MATLAB to conduct our MVPA analyses (Oosterhof et al., 2016).

##### Cued and distractor orientation

To examine whether there was information about the cued orientation, we trained a classifier to distinguish between the 4 possible cued orientations. These 4 orientations were divided into 2 pairs (see Figure 1B); 22.5° was paired with 112.5°, and 67.5° was paired with 157.5°. The classification of orientation was done within each pair separately, and the decoding accuracies were then averaged over pairs. This ensured that the cued orientation was orthogonal to the distractor orientation and comparison orientation. For example, in pair 1, cued orientation 22.5° and 112.5° occurred equally often with each of distractor orientations 67.5° and 157.5° (see Figure 1B). Cued and distractor orientations were also orthogonal to the rotation decision, so that only the orientation of the stimulus, and not the participant’s decision about it, could drive the classifier. Because the decoding was done within a pair, theoretical chance level was 50%. The decoding analysis for the distractor orientation was the same, as the possible cued and distractor orientations were identical.

##### Rotation direction

We used coding of the rotation direction as a measure for coding of the decision participants had to make. To determine whether there was information about the rotation direction, we trained a classifier to distinguish between a clockwise and anti-clockwise rotation. The rotation direction was orthogonal to both the cued and distractor orientation conditions (e.g., a clockwise decision was equally associated with all 4 possible orientations). There were 2 possible rotation directions, giving a theoretical chance level of 50%.

##### Response button

For completeness, we also examined coding of the response button. For this, we trained a classifier to distinguish between the two buttons participants used to indicate a ‘correct’ or ‘incorrect’ rotation shown on the response screen. We chose to decode the correct response button, that should have been pressed on each trial, rather than the pressed response button to maintain the full counterbalancing of trial types. We do not expect error trials to introduce much noise, given the high behavioural accuracy of mean 99.67% (after replacing error trials where possible). The correct response button was orthogonal to the rotation direction decision, and decoding of the response button is likely driven by motor preparation and execution.

#### 2.1.5 Channel searchlight

We performed an exploratory channel searchlight analysis to investigate which MEG sensors were driving the observed classification accuracies of 1) the cued orientation, 2) the distractor orientation, and 3) the rotation direction. We followed an established pipeline (Grootswagers et al., 2019; Robinson et al., 2019, 2022), where we ran the decoding described in section *2.1.5*. for a cluster of sensors around each MEG sensor. We used a Linear Discriminant Analysis instead of SVM classifier for this analysis to reduce computing time. To obtain the clusters, we took the closest neighbouring channels for each MEG channel, resulting in 2 to 7 neighbours per channel. We then ran the decoding for each sensor cluster and timepoint and stored the obtained decoding accuracies in the centre channel, resulting in a time-by-channel decoding accuracy map for each participant and decoding analysis. For visualisation purposes, we averaged the topographies across 200ms time bins, with the exception of the first topography (–100 to 0ms) and the last topography (2400 to 2500ms), which were averaged over 100ms time bins instead.

Previous work has shown that eye-movements can contribute to decoding, including decoding of orientation (Linde-Domingo & Spitzer, 2024; Mostert et al., 2018; Quax et al., 2019). In the present study, participants were instructed to fixate on a bullseye presented during the whole trial, and all stimuli were presented at fixation. In addition, the stimulus was presented for 150ms only, reducing the incentive to make saccades. However, we cannot fully rule out the contribution of eye-movements. The channel searchlight analysis may provide some insight, as one would expect the decoding of visual information to be driven by posterior brain regions, whereas any eye movement-related signals would likely come from frontal sensors near the eyes.

#### 2.1.6. Statistics

We used Bayesian statistics to determine the evidence for the coding of information under the null hypothesis (chance decoding) and the alternative hypothesis (above-chance decoding), for each time-point (Dienes, 2011; Jeffreys, 1998; Kass & Raftery, 1995; Rouder et al., 2009; Wagenmakers, 2007), using the Bayes Factor package in R (Morey & Rouder, 2018). To test whether the decoding accuracy was above chance, we used a half-Cauchy prior for the alternative, centred around d = 0, with the default width of 0.707 (Jeffreys, 1998; Rouder et al., 2009; Wetzels et al., 2011). To exclude irrelevant effect sizes, we excluded the d = 0 to 0.5 interval from the prior (Morey & Rouder, 2011; Teichmann et al., 2022), and used a point null at d = 0. To test whether there was stronger information coding for the cued compared to the distractor orientation, we calculated the difference between decoding accuracies for the cued and distractor orientation. We used a half-Cauchy prior, centred around 0, with the same width and null interval described above, for the alternative hypothesis to capture directional effects (cued > distractor).

Bayes Factors (BFs) of <1 show evidence for the null hypothesis, while BFs of >1 show evidence for the alternative hypothesis. BFs between 1/3 and 3 are typically interpreted as showing insufficient evidence, BFs < 1/3 or BFs >3 as substantial evidence, and BFs < 1/10 or BFs >10 are interpreted as strong evidence (Wetzels et al., 2011). In this study, we defined the onset of strong evidence for above-chance decoding as the second consecutive timepoint with a BF >10, and the return to baseline decoding as the second consecutive timepoint with a BF<1/10.

### 2.2. fMRI acquisition and analysis

#### 2.2.1. Participants

The second part of the study consisted of a behavioural training session and an fMRI session carried out at the MRC Cognition and Brain Sciences Unit (Cambridge, UK). 42 healthy volunteers participated in the initial training session. Of these, 27 volunteers performed well enough to participate in the fMRI session (at least 90% accuracy in the final run of the training session). The fMRI data from 3 participants were excluded due to excessive movement, failure to complete the experiment, or low task performance (accuracy < 80% in the fMRI session), resulting in a final fMRI sample of 24 participants (15 female/9 male, 23 righthanded/1 lefthanded, mean age = 27.33 years, SD = 5.53). One of these participants also participated in the MEG experiment and 23 of the participants were new recruits. Participants received £6 for participation in the training session and £20 for participation in the fMRI session, as well as £2.50 – £3 travel costs per session. All participants provided informed consent, and the study was approved by the Cambridge Psychology Research Ethics Committee.

#### 2.2.2. Stimuli and experimental procedure

##### fMRI session

Participants performed the same task as in the MEG, except for the following 3 changes due to the lower temporal resolution of fMRI. First, the order of consecutive trial types was not fully randomised as it was in the MEG version, but was instead counterbalanced within each run, making sure each combination of cued orientation x distractor orientation x comparison orientation was equally likely to follow each other combination as well as itself. The order of the rotation direction was balanced separately within each run. Secondly, to keep the order of consecutive trial types intact, error trials were not repeated. Thirdly, participants did not receive trial-wise feedback to avoid feedback of the previous trial influencing the signal of the next trial. Participants still received feedback about their accuracy at the end of each block. Participants completed 8 runs, with 4 blocks per run, and 32 trials per block. The total number of blocks, and the number of trials per block stayed the same as the MEG experiment, however, the blocks were divided into 8 runs of 4 blocks, instead of 16 runs of 2 blocks. One participant completed 7 instead of 8 runs.

The timing of the task was kept the same at the MEG experiment. Due to the lower temporal resolution of fMRI, we are not able to separate the effect of attention from the decision-making processes in time. However, the rotation direction, about which participants made a decision, was an orthogonal dimension to both the cued and distractor orientation. This means that we can separate the effect of attention on stimulus coding from the decoding of the decision information.

##### Training session

The training session was the same as described in the MEG experiment section.

#### 2.2.3. fMRI acquisition

fMRI scans were acquired using a Siemens 3 Tesla Prisma-Fit scanner (Siemens Healthcare, Erlangen, Germany), with a 32-channel head coil at the MRC Cognition and Brain Sciences Unit, Cambridge, UK. We used a multi-band T2*-weighted echo planar imaging (EPI) acquisition sequence with the following parameters: repetition time (TR) = 1208ms; echo time (TE) = 30ms; flip angle = 67 degrees, field of view = 192 mm, multi-band acceleration factor = 2, no in-plane acceleration, in-plane resolution = 3 x 3 mm, 38 interleaved slices of 3 mm slice thickness with 10% interslice gap. T1-weighted MPRAGE structural images were acquired at the start of the session (resolution = 1 x 1 x 1 mm). The stimuli were presented on a NNL LCD screen (resolution = 1920 x 1080, refresh rate = 60 Hz) using the Psychtoolbox extension in MATLAB (Brainard, 1997; Kleiner et al., 2007; Pelli, 1997). Participants used a 2-button response pad to respond on each trial. To get comfortable with performing the task in the MRI scanner, participants completed 2 practice blocks of the task during the acquisition of the structural scan at the start of the scanning session. Feedback about accuracy was given at the end of each of these practice blocks.

#### 2.2.4. Pre-processing and first-level model

The data were pre-processed using SPM 8 (Wellcome Department of Imaging Neuroscience) in MATLAB. EPI images were converted to NIFTII format, spatially realigned to the first image and slice-time-corrected (slice timing correction used the routine from SPM12 to allow for multiband acquisitions). Structural scans were co-registered to the mean EPI image and normalised to derive the normalisation parameters needed for the definition of the regions of interest (ROIs).

To obtain activation patterns for the MVPA analysis, we estimated a General Linear Model (GLM) for each participant. There were 16 regressors per run, reflecting a combination of 4 cued orientations x 2 distractor orientations x 2 rotation directions (see Figure 1B). Whole trials were modelled as a single events, lasting from the onset of the first stimulus until the response was given, to account for trial-to-trial variability in response time (Woolgar et al., 2014). Regressors were convolved with the hemodynamic response function of SPM8.

#### 2.2.5. Regions of interest

13 frontal and parietal MD ROIs were taken from the parcellated map provided by Fedorenko et al (2013), which is freely available online at imaging.mrc-cbu.cam.ac.uk/imaging/MDsystem. The definition of the MD network is activation based, as the map indexes regions that show a univariate increase in activation with increased task demands, across a range of tasks. This is an updated definition of the MD system, with a high degree of overlap with the previous definition, derived from meta-analytic data (Duncan & Owen, 2000) that we used in previous work (Jackson et al., 2016; Jackson & Woolgar, 2018; Woolgar, Williams, et al., 2015; Woolgar, Afshar, et al., 2015), and a more recent definition derived from multimodal imaging (Assem et al., 2020). The ROIs comprised left and right anterior inferior frontal sulcus (aIFS; center of mass (COM) = ±35 47 19, volume = 5.0 cm^3^), left and right posterior inferior frontal sulcus (pIFS; COM ±40 32 27, 5.7 cm^3^), left and right premotor cortex (PM; COM ±28 −2 56, 9.0 cm^3^), left and right inferior frontal junction (IFJ; COM ±44 4 32, 10.1 cm^3^), left and right anterior insula/frontal operculum (AI/FO; COM ±34 19 2, 7.9 cm^3^), left and right intraparietal sulcus (IPS; COM ±29 −56 46, 34.0 cm^3^), and bilateral anterior cingulate cortex (ACC; COM 0 15 46, 18.6 cm^3^). Early visual cortex was defined as area BA 17 (center of mass = −13, −81, 3/16, −79, 3, volume = 54 cm^3^) from the Brodmann template provided with MRIcro (Rorden & Brett, 2000). ROIs were deformed into native space by applying the inverse of the normalisation parameters for each participant.

#### 2.2.6. Decoding analysis

We used the same decoding analysis as described in the MEG experiment but in the MEG experiment, we used MVPA to determine what information is present in the pattern of activation for each *point in time*, whereas for the fMRI data, we determine what information is present in the pattern of activation *for each ROI*. Within each ROI, we trained an SVM classifier on the betas obtained from the GLM to distinguish the conditions of interest. The classifier performance was determined by using a leave-one-run-out cross-validation. Decoding accuracies were then averaged over hemispheres.

#### 2.2.7. Statistics

To test for an effect of attention, MD region, and the interaction between these factors, we used a Bayesian analysis of variance (ANOVA) with attention (cued vs. distractor orientation) and MD region (aIFS, pIFS, PM, IFJ, AI/FO, IPS, and ACC; data collapsed across hemispheres) as within-subject factors, using the default Jeffries prior of medium width (1/2) (Rouder et al., 2012). We conducted further Bayesian t-tests using the same parameters described in the MEG experiment. To determine the contribution of individual MD regions and V1 to the effect of attention on stimulus coding, we performed Bayesian tests of the difference between decoding of the cued and distractor orientation per MD ROI and V1. In addition, we performed Bayesian tests for the decoding of the cued orientation, distractor orientation, and rotation direction against chance for the mean MD regions, per individual MD region and for V1. We used the same parameters for these tests as described in the MEG section (2.1.6).

### 2.3. Model-based MEG-fMRI fusion

To gain insight into the coding of attended information, unattended information, and decision-related information in both space and time simultaneously, we used model-based MEG-fMRI fusion (Cichy et al., 2014, 2016; Hebart et al., 2018). This method uses Representational Dissimilarity Matrices (RDMs) (Kriegeskorte et al., 2006, 2008) to abstract away from the imaging modality, allowing us to compare the pattern similarity across neuroimaging modalities. In addition, we can abstract away from the pattern of activation of specific participants, comparing the representational similarity across different groups. Model-based MEG-fMRI fusion further allows us to determine the match between the representational structure for each timepoint (MEG), and each ROI (fMRI), which can be uniquely explained by a theoretical model (Hebart et al., 2018) (see Figure 6A for an overview of this method). We used 3 orthogonal models, coding for the *cued orientation*, *distractor orientation*, and *rotation direction*.

The RDMs consisted of a 16 by 16 matrix comprising each combination of cued orientation (4) x distractor orientation (2) x rotation direction (2) (see Figure 1B for an overview of combinations). For each cell in the RDM, we used decoding accuracy as our measure of dissimilarity, with greater decoding accuracy reflecting greater dissimilarity of activation patterns (i.e., distinctiveness of patterns between the conditions). Decoding accuracies were obtained for each cell in the RDM (i.e., pair of trial types) using the same cross-validation procedure as described above and were z-scored per ROI and timepoint. For the MEG data, we constructed an RDM at each timepoint. For each cell in the RDM, we obtained decoding accuracies using a 25ms sliding window centred around each timepoint. For the fMRI data, we constructed an RDM for 9 different ROIs: the mean MD, V1, aIFS, pIFS, PM, IFJ, AI/FO, IPS, and ACC. The mean MD ROI was obtained by calculating the RDM for each individual MD region and subsequently averaging RDMs across regions^1^. The individual MD ROIs, except for the ACC, were obtained by calculating separate RDMs for the left and right hemispheres, and then averaging across RDMs. We did this separately for each participant, and then averaged the RDMs over participants, resulting in a single RDM per timepoint for MEG, and for each of the 9 ROIs for fMRI.

We constructed 3 model RDMs: for the cued orientation, distractor orientation, and rotation direction (see Figure 6A). The cued and distractor orientation models coded for: the same orientation (0), a difference of 45° (0.5), or a difference of 90° (1); the rotation direction model coded for the same rotation (0) or a different rotation (1).

We used commonality analysis to estimate the shared variance between 3 RDMs: the MEG RDM for each point in time, the fMRI RDM for each ROI, and the model RDM. For each timepoint, ROI, and model, we calculated the difference between two squared semi-partial correlation coefficients, using Spearman correlation. Both semi-partial correlation coefficients reflected the proportion of variance shared between MEG and fMRI; one with *all models* partialled out from the MEG RDM, and the other with *all models except the model of interest* partialled out from the MEG RDM. By comparing the variance when our model of interest was included or not, we get a measure of the variance shared between MEG and fMRI that can be uniquely explained by each model (Hebart et al., 2018). In this way we derive a time-course of the model fit for each region.

#### 2.3.1. Statistics

Because we averaged the RDMs over participants for both MEG and fMRI, we had a single RDM for each timepoint for MEG, and a single RDM for each ROI for fMRI. It was therefore not possible to calculate random effects statistics at the group level, as was done for the MEG and fMRI decoding analyses. We therefore used a permutation test. To estimate the null distribution, we computed 10,000 permutations by shuffling the rows and columns of the group average MEG RDM and ran model-based MEG-fMRI fusion for each ROI using the permuted MEG matrices. For each ROI, we then determined a cluster-inducing threshold for each timepoint by choosing the 95^th^ percentile. We determined the largest maximum cluster size in the null distribution. We did this over both ROIs to account for multiple comparisons over our 2 ROIs. We compared the clusters in our fusion data to the maximum cluster size obtained from the null distribution (equivalent to p<0.05, one-tailed, corrected for multiple comparisons for the two ROIs and all the timepoints).

## 3. RESULTS

### 3.1. MEG results

#### 3.1.1. Behavioural results for MEG session

Participants performed with high accuracy in the MEG session. The mean accuracy was 94.23% (SD = 3.90) before replacing error trials (see Methods), and 99.67% (SD = 0.56) after replacing error trials where possible, which means very few error trials were included in the analysis (0.33% of trials on average). The mean response time was 667ms (SD = 104ms).

#### 3.1.2. Coding of the cued and distractor orientation over time

To examine the timepoints at which attended visual information was preferentially coded over unattended visual information, we compared the decoding accuracies for the same orientations when presented as the cued or distractor orientation for each point in time. Critically, the coding of the cued and distractor information could not be driven by decision-related information, as this was an orthogonal dimension in the design. The decoding accuracies for the cued and distractor orientations are shown in Figure 2, the topographies below the plots show the contribution of individual sensors to the decoding accuracy. There was strong evidence for above-chance orientation decoding for the cued orientation starting at 85ms after stimulus onset, which was maintained until after the mean response time. The distractor stimulus orientation could be decoded with a very similar onset to the cued orientation, with strong evidence for above-chance coding from 90ms after stimulus onset. However, unlike the coding of the cued information, coding of the distractor orientation was not sustained over time, dropping back to chance within 420ms after stimulus onset. The difference between coding of the cued and distractor orientations, which reflects the attentional selection and maintenance of task-relevant information, was evident from 215ms after stimulus onset. This effect of attention, with stronger coding of the cued compared to distractor orientation, was maintained until after the mean response time.

**Figure 2.**
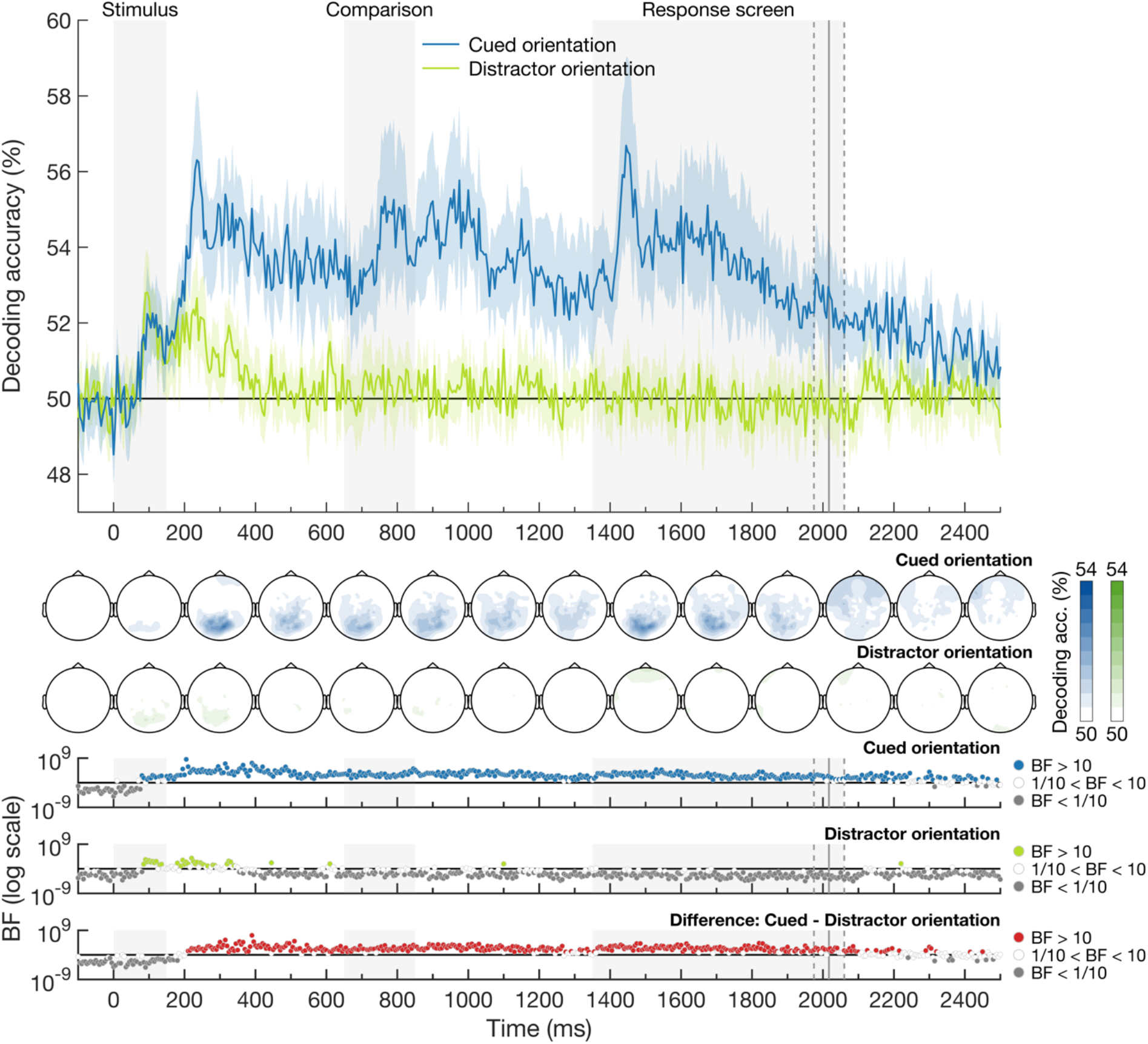
Decoding accuracy of the cued and distractor orientation over time. Classifiers were trained to discriminate stimulus orientation when the orientation was either cued (blue) or not (light green). Theoretical chance is 50% decoding accuracy, and shaded areas around the plot lines show the 95% confidence intervals. The grey shaded areas show from left to right: 1) when the stimulus was on the screen (oriented lines in cued and distractor colours; labelled Stimulus), 2) when the comparison line was on the screen (labelled Comparison), and 3) when the response screen was shown (labelled Response screen). The response screen remained on until participants responded (or 3 secs). The vertical grey lines show the mean response time with 95% confidence intervals. The topographies below the plot show the sensor searchlight decoding results for the cued orientation (top, blue) and distractor orientation (bottom, green). The topographies were averaged over 200 ms time bins, except for first topography (–100 to 0ms) and the last topography (2400 to 2500ms) which are based on a 100ms time bin. The colour bars next to the topographies show the scale of the decoding accuracies. There do not seem to be specific frontal contributions to the orientation decoding accuracy, suggesting that eye-movements do not have a strong contribution to the observed orientation decoding. Bayes factors are given below the plot on a logarithmic scale for the cued orientation (blue), the distractor orientation (light green) and the difference between the cued and distractor orientation (red). BFs below 1/10 are shown in grey, indicating strong evidence for the null hypothesis. BFs above 10 are shown in the plot colour 511 (blue for the cued orientation, light green for the distractor orientation, and red for the effect of attention), indicating strong evidence for the alternative hypothesis. BF between 1/10 and 10 are shown in white. There was strong evidence for coding of the cued orientation from 85ms after the onset of the stimulus which was maintained until after the response was given. There was strong evidence for coding of the distractor orientation from 90ms after the onset of the stimulus, but this was not sustained over time. There was strong evidence for an effect of attention from 215ms after the onset of the stimulus until after the mean response time.

#### 3.1.3. Coding of the rotation direction over time

To examine the time-course of decision-related information coding, we determined the coding of the rotation direction over time by training a linear classifier to distinguish between clockwise and anti-clockwise rotations. Participants had to actively manipulate the cued orientation in combination with the comparison orientation to determine the rotation. The decoding of the rotation direction over time is shown in Figure 3. We observed above-chance decoding of the rotation direction from 170ms after the onset of the comparison line, 820ms after the onset of the attended and unattended orientations, which was maintained until after the mean response time. For completeness, we also decoded the correct response button (Figure 3). We observed information about the response button from 1580ms onwards, 230ms after the onset of the response screen.

**Figure 3.**
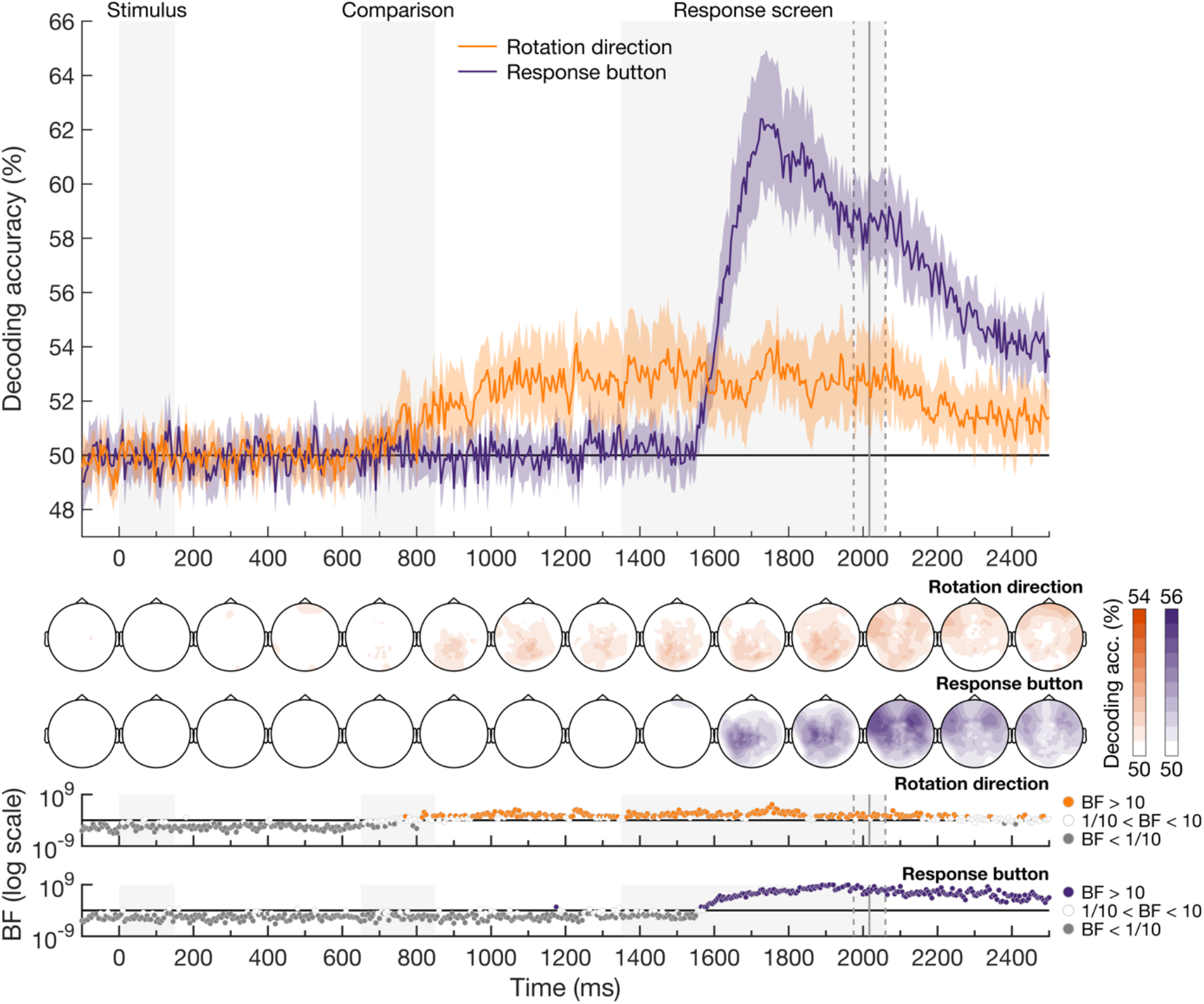
Decoding accuracy of the rotation direction and the correct response button over time. Classifiers were trained to discriminate the rotation direction, which could be clockwise or anti-clockwise. The rotation direction (orange), about which participants had to make a decision, could be decoded from 170ms after the onset of the comparison line (820ms after stimulus onset) and was maintained until after the mean response time. The time-course of decoding accuracies for the response button (purple) was included for completeness. The topography for the decoding of the rotation direction is shown below the plot. Plotting conventions are as in Figure 2.

### 3.2. fMRI results

#### 3.2.1. Behavioural results for fMRI session

Participants performed the task in the fMRI session with high accuracy (mean accuracy = 94.45%, SD = 4.91). Included participants all had a mean accuracy above 80%. The mean response time was 658ms (SD = 140ms) from response screen onset.

#### 3.2.2. Coding of the cued and distractor orientation in the MD regions and V1

To examine whether the effect of attention observed in the MEG data pertained to V1 and the MD regions, we decoded the cued and distractor orientations for these ROIs. We were able to separate these conditions due to the orthogonal design. The decoding accuracies for the cued and distractor orientations, averaged across the MD regions and in V1, are shown in Figure 4A. We used a Bayesian ANOVA with attention (orientation coding of cued vs. distractor stimuli) and MD region (aIFS, pIFS, PM, IFJ, AI/FO, IPS, and ACC; data collapsed across hemispheres) as within-subject factors. We found a main effect of attention (BF > 100), no main effect of region (BF = 0.44), and an interaction between attention and region (BF = 9.60), justifying a further analysis of the effect of attention in individual MD regions (see Figure 4B). There was strong evidence for an effect of attention (cued > distractor coding) in aIFS, pIFS, PM, IFJ, and IPS, and some evidence for an effect of attention in AI/FO and ACC (see Figure 4B).

**Figure 4.**
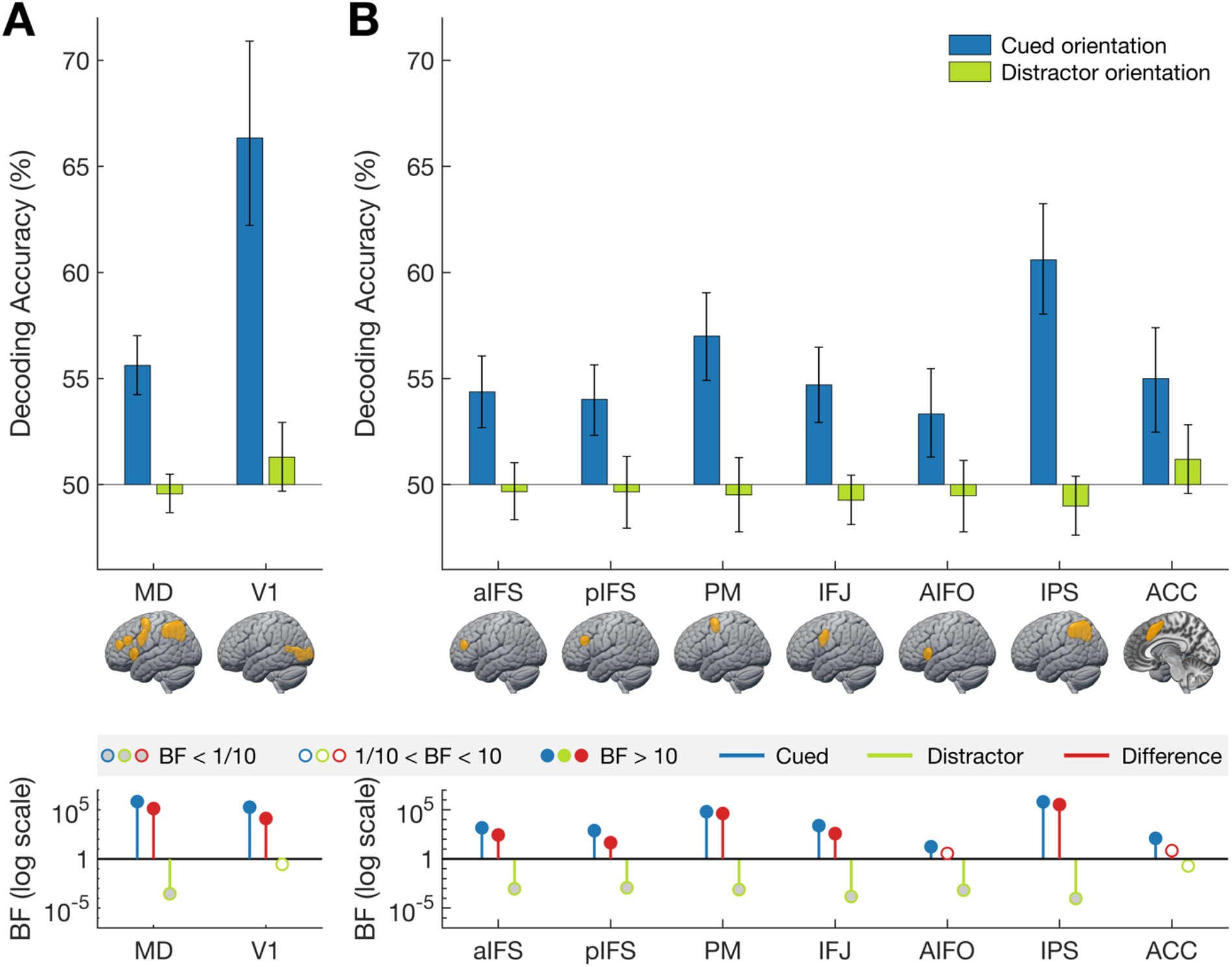
Decoding accuracy of the cued and distractor orientation (A) averaged across the MD regions and in V1; and (B) in individual MD regions. In both **A** and **B**, classifiers were trained to discriminate stimulus orientation when the orientation was either cued (blue) or not (light green). Theoretical chance is 50%, and error bars indicate 95% confidence intervals. The Bayes factors for the decoding accuracies are shown below the plot in blue (cued orientation) and light green (distractor orientation), and the BFs for the difference (cued>distractor coding) are shown in red. All BFs are shown on a logarithmic scale. BFs > 10 are marked in the plot colour, BFs < 1/10 are marked in grey, and BFs in between this range are marked in white. In all of the MD regions, and in V1, there was strong evidence for coding of the cued orientation, but not for the distractor orientation. There was strong evidence for an effect of attention for most of the MD regions (aIFS, pIFS, PM, IFJ, AI/FO, and IPS) as well as average MD and V1.

To determine whether the orientation coding for the cued and distractor orientations was above chance, we performed Bayesian t-tests for the mean MD regions as well as the individual MD regions. There was strong evidence for above-chance coding for the cued orientation in the mean MD regions (mean accuracy = 55.62%, BF > 100), while decoding of the distractor orientation was at chance (mean accuracy = 49.56%, BF < 0.01). There was above-chance decoding for the cued orientation in all of the MD regions individually. Strong evidence for chance level decoding for the distractor orientation was found for aIFS, pIFS, PM, IFJ, AI/FO, and IPS (BF < 0.01). There was evidence for chance decoding for the distractor orientation in ACC (BF = 0.20, and the 95% confidence intervals for this region overlapped with chance, consistent with evidence for the null.

We also found an effect of attention in V1 (BF > 100), with stronger coding for the cued orientations compared to the distractor orientations. V1 coded for the cued orientation (mean accuracy = 66.34%, BF>100), while when these identical orientations were distractors there was evidence for chance decoding (mean accuracy = 51.29%, BF = 0.28), and the 95% confidence intervals overlapped with chance, consistent with evidence for the null.

#### 3.2.3. Coding of the rotation direction in the MD regions and V1

To examine whether there was information about the rotation direction, which is our indicator of the decision, in the MD regions and V1, we trained a linear classifier to distinguish between a clockwise and anti-clockwise rotation direction. Since error trials were included in the analysis to maintain the orthogonality of the different features, on a small number of trials (∼6%), the decoded rotation direction does not reflect the decision of the participant, but the predominant influence should be the decision. Figure 5A shows the decoding accuracies for the rotation direction in the MD regions and V1. We used Bayesian t-tests to determine the evidence for chance versus above-chance decoding. We observed strong evidence for above-chance coding of rotation direction across the MD regions on average (mean accuracy = 52.43%, BF = 45.89), and in PM and IPS individually (Figure 5B). Although the average decoding accuracy found in V1 was similar to the MD regions, greater variance in V1 was reflected in an inconclusive BF (mean accuracy = 52.67%, BF = 1.85). There was substantial, but not strong, evidence for chance decoding in aIFS, pIFS, and IFJ, and insufficient evidence for rotation direction coding in the other MD regions.

**Figure 5.**
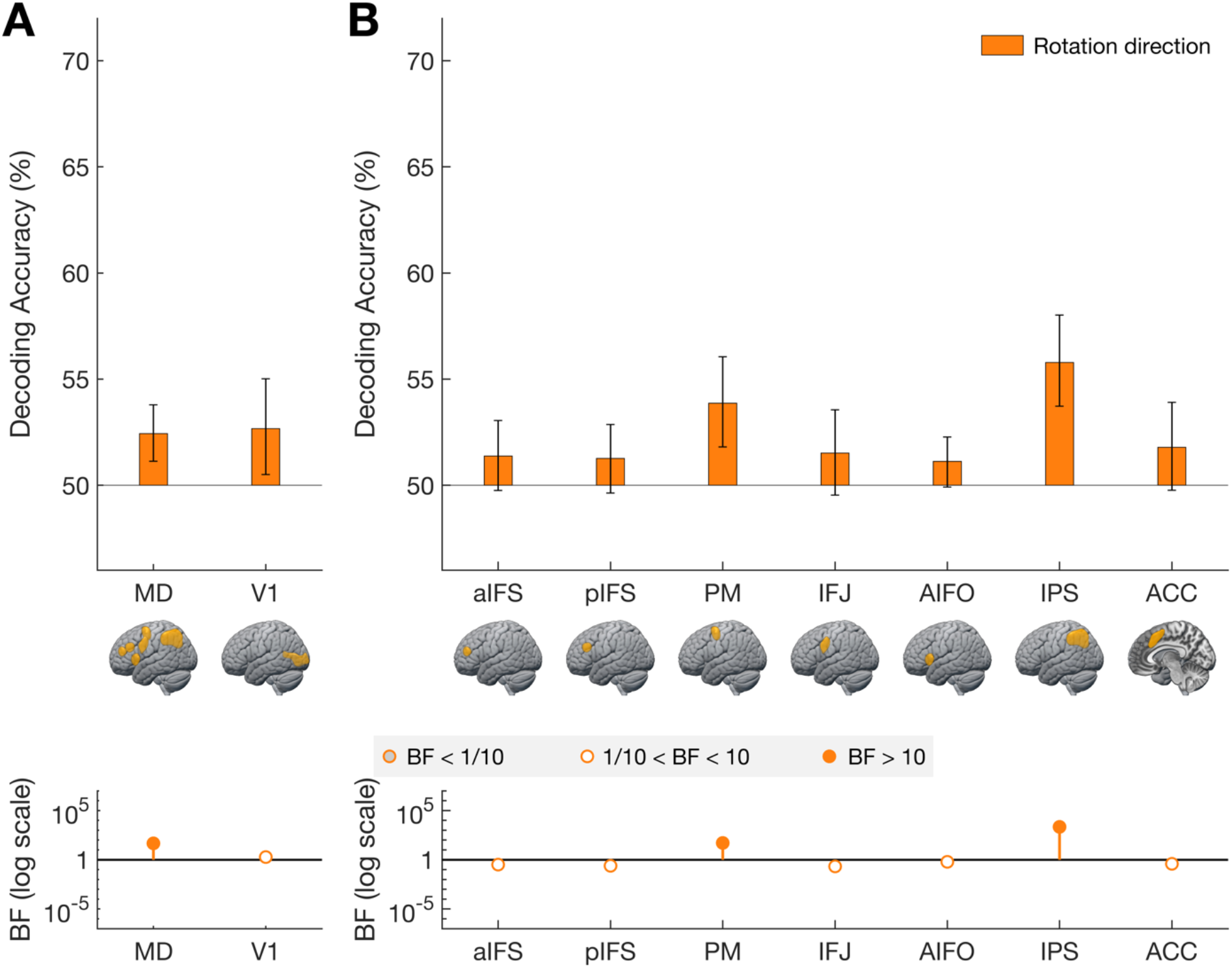
Decoding accuracy of the rotation direction (decision) (A) averaged across the MD regions and in V1 and (B) in individual MD regions. In both **A** and **B**, classifiers were trained to discriminate the rotation direction, which could be clockwise or anti-clockwise. Plotting conventions are as in Figure 4. The rotation direction could be decoded from the MD regions (average, PM and IPS), while there was insufficient information for rotation direction coding in V1.

### 3.3. Model-based MEG-fMRI fusion results

We used model-based MEG-fMRI fusion to ask whether the MD regions coded attended information *before* a decision was made. The commonality coefficient shown in Figure 6A refers to the part of the variance that is shared between the MEG and fMRI RDMs which can be uniquely explained by our models for the cued orientation, distractor orientation, and rotation direction. In other words, it plots the dynamics with which the MD pattern, observed in fMRI and explained by each of our theoretical models, arises in the MEG data. We ran this analysis for the mean MD regions, individual MD regions, and V1.

**Figure 6.**
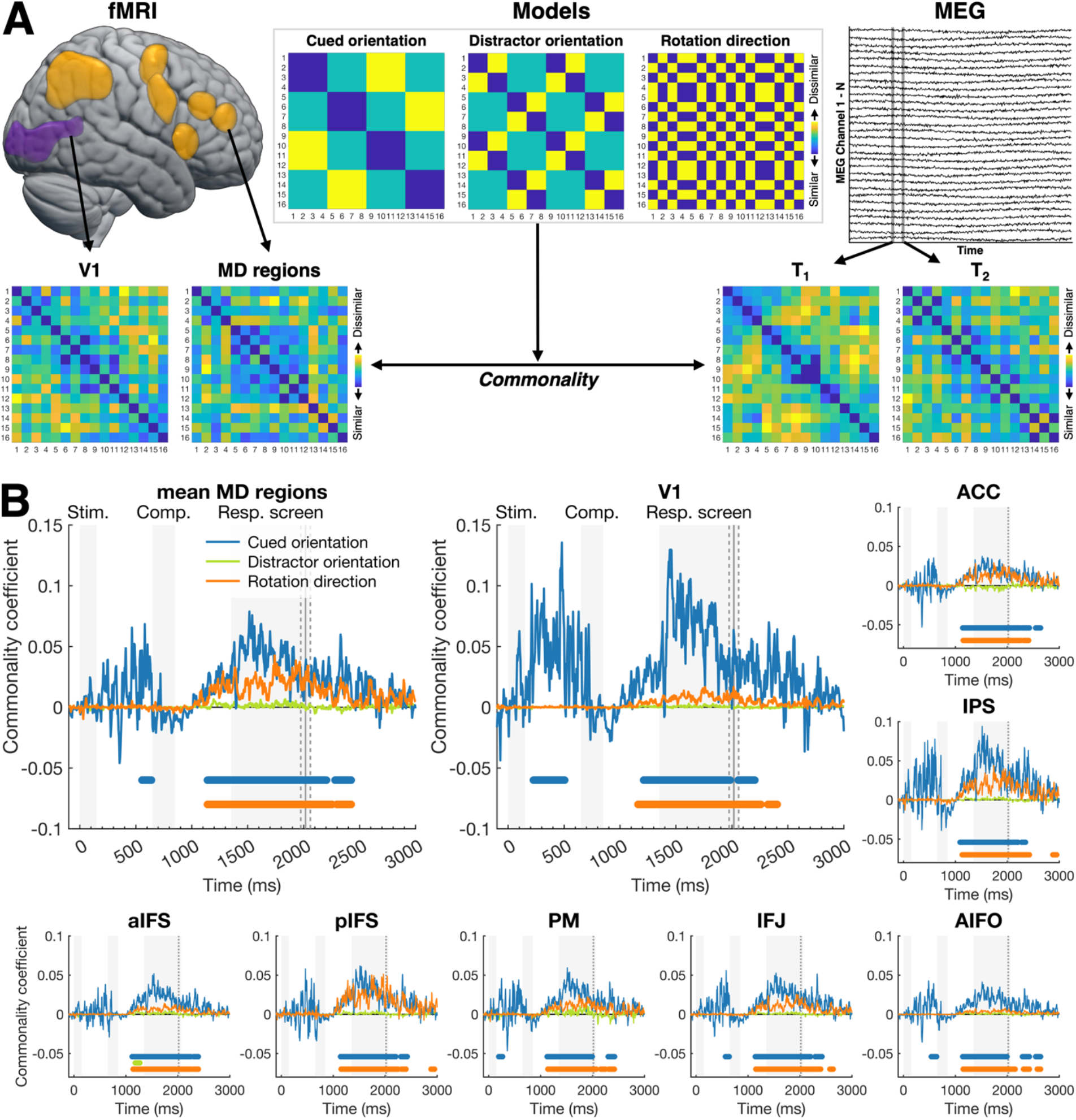
Model-based MEG – fMRI fusion methods (A) and results over time (B). **A** gives an overview of the model-based MEG – fMRI fusion method. We used RDMs to abstract away from the imaging modality. Each cell in the RDM corresponds to the dissimilarity between 2 specific trial types. A trial type is a combination of the cued orientation, distractor orientation, and rotation direction. Trial types 1-16 in the RDMs correspond to the 16 trial types listed in Figure 1B. We created an RDM for each region of interest (V1 and mean MD regions) for the fMRI data, for each time-point in the MEG data, and for the 3 models (cued orientation, distractor orientation, and rotation direction). We then determined commonality: the part of the variance that is shared between the MEG RDM for each timepoint and the fMRI RDM for each ROI, that can be uniquely explained by each theoretical model. This results in a time-course of commonality coefficients for each ROI and model. Note that the ROIs are plotted by projecting a 3D region onto the cortical surface. **B** show the time-course of commonality all ROIs: the mean MD regions, V1, aIFS, pIFS, PM, IFJ, AI/FO, IPS, and ACC respectively. The different lines depict the commonality explained by the model for the cued orientation (blue), the distractor orientation (light green), and the rotation direction (orange). Significant timepoints (equivalent to p<0.05, one-tailed) were obtained with a cluster-corrected randomisation test, corrected for multiple comparisons across ROIs and timepoints, and are shown at the bottom of the plots. Plotting conventions are as in Figure 2 and 3 except that the dots in the bottom of this plot show significant time-points, not Bayes Factors.

#### 3.3.1. Cued orientation

In the MD regions, there was a significant cluster of commonality for the cued orientation from 555ms until 640ms after the onset of the stimulus (Figure 6B). Critically, this cluster occurred *before* the onset of the comparison line, and therefore before participants could make their decision. A second cluster started at 1140ms, 490ms after the onset of the comparison line and was maintained until after the average response time. The observed pattern was similar for all individual MD regions. However, the initial cluster before the onset of the comparison line only reached significance for PM, IFJ, and AI/FO. In V1, cued orientation commonality coefficients were significant from 225ms until 505ms after the onset of the stimulus, and again from 1210ms, 560ms after the onset of the comparison line, until after the mean response time.

#### 3.3.2. Distractor orientation

There were no significant clusters of commonality for the distractor orientation in the mean MD region, V1, or in the individual MD regions except for aIFS. In aIFS, a brief cluster was observed between 1180 and 1270ms after stimulus onset.

#### 3.3.3. Rotation direction

We observed a cluster of significant commonality for the rotation direction, which is an index of the decision participants had to make, in the mean MD regions starting from 1140ms, 490ms after the onset of the comparison line, which lasted until after the participant responded. Individual MD regions showed a similar pattern. V1 also showed a similar pattern over time, with a significant rotation direction cluster from 1160ms, 510ms after comparison line onset.

## 4. DISCUSSION

A wealth of fMRI, M/EEG and non-human primate studies have shown preferential coding of cued compared to distractor information (Barnes et al., 2022; Battistoni et al., 2020; Bode & Haynes, 2009; Goddard et al., 2022; Grootswagers et al., 2021; Harel et al., 2014; Jackson et al., 2016; Kaiser et al., 2016; Li et al., 2007; Moerel et al., 2022; Rao et al., 1997; Sigala et al., 2008; Stiers et al., 2010; Stokes et al., 2013; Waskom et al., 2014; Woolgar, Afshar, et al., 2015; Woolgar, Williams, et al., 2015). However, most paradigms manipulate attention by requiring participants to respond to the cued (but not the distractor) stimulus, which means they cannot distinguish between preferential coding of cued information due to attentional selection and maintenance of relevant information, and coding due to decision-making processes performed on the attended information. We used MEG, fMRI, and model-based MEG-fMRI fusion to distinguish between these explanations. Our MEG data showed that attention affects stimulus processing around 215ms after stimulus onset, before participants had the necessary information to make an explicit decision in our task. Then, when the comparison line was presented, the decision about the rotation direction could be decoded approximately 170ms later. Our fMRI data showed the MD regions coded attended visual information, and did so more strongly than distracting visual information, even though the decision made was orthogonal to the stimulus itself. The orthogonal decision representation was also present in the MD system, specifically in PM and IPS. We also found an effect of attention in V1, with stronger coding of the cued compared to distractor orientations, in line with previous studies (Jackson et al., 2016; Jehee et al., 2012; Pratte et al., 2013). Our model-based MEG-fMRI fusion analysis showed evidence for the coding of attended orientation information in the MD regions *before* participants could make a decision. These data are consistent with the interpretation of previous results as effects of attention on the processing of visual stimuli, with a stronger coding of attended compared to unattended visual information in the brain. This could be driven by an enhancement of the attended visual information, a suppression of the unattended visual information, or both.

The finding that the MD regions preferentially code for attended information over unattended information, even when the decision is orthogonal, is in line with the finding of Hon and colleagues (2006), who used univariate analyses to show that the average MD response reflected changes in cued information when no decision was needed, or when the decision was unrelated to the cued information. Our results take the inference beyond simple activation changes and confirm that decision-making processes are not required for the prioritisation of cued over distractor information.

Our results also show that, in addition to coding the attended information, MD regions also code for decision-related information. The PM and IPS, which are part of the MD network, held information about the rotation direction, which was the decision participants had to make. The time-course with which this information emerged, from about 170ms after the onset of the comparison line, fits well with a previous study which found a perceptual decision on a pure noise image could be decoded around 140-180ms (Bode et al., 2012). The fast time-course suggests the coding of rotation direction information could reflect the accumulation of decision-related information before the participant is able to respond. This is in line with the description of decision-making as the accumulation of evidence towards one of two decision alternatives until a decision boundary is reached (Gold & Shadlen, 2007; Heekeren et al., 2008; Ratcliff & McKoon, 2007). We cannot comment on the exact delay between the onset of decision information coding and the response, as our task involved a delayed response – participants could only generate a response (which was again orthogonal to the decision) after the response screen came on.

In addition to separating the effects of attention and decision-making, our study adds to the literature by combining neuroimaging techniques, giving us resolution in both space and time for a single paradigm. Our MEG data show that the effect of attention emerges *before* participants have all of the information needed to make a decision, while our fMRI data show that the MD regions code both attended visual information and the decision participants have to make about the stimulus. Moreover, we formally related the two results using model-based MEG-fMRI fusion in order to estimate the time-course of response in the MD network and V1 separately. The data suggest that visual information arises first in V1, but that both V1 and the MD regions code for information about the attended orientation *before* participants can begin to make a decision. Orthogonal decision-related signals in MD cortex arose only after the comparison line was presented. The MEG-fMRI fusion method is an exciting step towards resolving neural processes in both time and space, but is limited in sensitivity since it is only able to pick up effects that are present in both the MEG data and the fMRI data (see Cichy & Oliva (2020) for an overview of possible limitations). For instance, although our MEG data revealed transient coding of unattended information, this was not evident in the fusion results, presumably because the much slower time resolution of fMRI obscured its detection. This highlights the utility of considering the data from each neuroimaging technique separately, as well as together.

Our findings emphasise the adaptable response of the MD regions, in line with the literature in non-human primates showing the same neural populations can simultaneously encode multiple task parameters (Aoi et al., 2020; Fusi et al., 2016; Rigotti et al., 2013). For example, Aoi and colleagues (2020) show that a single neural population in monkey prefrontal cortex can maintain information about relevant and irrelevant sensory information, as well as the saccade response, over the course of a trial. Another line of evidence for the flexible response of the MD regions comes from fMRI studies in humans. These studies show that multivariate patterns of activity code for a range of information in different tasks (Woolgar et al., 2016), with single MD voxels re-used to code multiple task features between tasks (Jackson & Woolgar, 2018). In addition, Woolgar and colleagues (2015) found stronger visual object coding in the MD regions for cued relative to distractor objects, and Jackson and colleagues (2016) reported stronger MD coding of cued compared to distractor features of the same object. Moreover, MD coding of cued information can be selectively impaired by non-invasive stimulation of the right dorsolateral prefrontal cortex, one of the nodes in the MD network (Jackson et al., 2021). Our study also provides evidence for a multi-faceted response of the MD regions, although we cannot determine whether this is driven by the same population of neurons within those brain areas. The multifaceted MD response observed in our study, reflecting relevant visual information as well as decisions about this information, is consistent with the proposal that these regions play a key role in associating different types of information by integrating information over multiple sources (Duncan, et al., 2020). The MD system is well placed to achieve this, being widely distributed and strongly interconnected across the cortex (Assem et al., 2020; Duncan et al., 2020). In addition, our results emphasise an explicit role of the MD regions in selective attention, with preferential coding of the information that is task relevant separately from functional decisions.

In summary, our MEG results show that attention affects the representation of visual stimuli in advance of decision-making processes, while the fMRI results show that the MD regions code for attended visual information as well as decisions about this information. The MEG-fMRI fusion results show the spatio-temporal unfolding of these processes, suggesting that the MD regions represent information about the attended stimulus before participants can begin to make a decision. These results emphasise a key role for the MD regions in selective attention and are consistent with the proposal that these regions integrate different types of information for decision-making. Our multimodal data demonstrate that selective attention and decision-making have separable bases in neural coding.

## ACKNOWLEDGEMENTS

This work was supported by the Australian Research Council (ARC) Centre of Excellence in Cognition and its Disorders (CE110001021), International Research Training Program Scholarships from Macquarie University awarded to DM, an ARC Discovery Project (DP170101840) awarded to ANR and AW, and by the Medical Research Council (UK) intramural funding (SUAG/093/G116768) awarded to AW. The authors acknowledge the facilities and scientific and technical assistance of the National Imaging Facility, a National Collaborative Research Infrastructure Strategy (NCRIS) capability, at Macquarie University. For the purpose of open access, the authors have applied a Creative Commons Attribution (CC BY) licence to any Author Accepted Manuscript version arising from this submission.

## Footnotes

1 We could instead have calculated the RDM by extracting the data from one large ROI including all MD regions; in practice this gave a similar RDM so was not considered further.

